# Targeted proteomics of postmortem human brain reveals neurobiologic heterogeneity in Alzheimer’s disease using NULISA technology

**DOI:** 10.64898/2026.07.01.735558

**Authors:** Sara Rose Dunlop, Sarah J. Lincoln, Zhongwei Peng, Neill R. Graff-Radford, Christian Lachner, Gregory S. Day, Jessica F. Tranovich, R. Ross Reichard, Dennis W. Dickson, Ronald C. Petersen, Bradley F. Boeve, Aivi Nguyen, Lea. T. Grinberg, Jonathan Graff-Radford, Alicia Algeciras-Schimnich, Melissa E. Murray

**Affiliations:** Department of Neuroscience, Mayo Clinic Jacksonville, Florida USA; Department of Quantitative Health Sciences, Mayo Clinic Jacksonville, Florida USA; Department of Neurology, Mayo Clinic Jacksonville, Florida USA; Department of Laboratory Medicine and Pathology, Mayo Clinic Rochester, Minnesota USA; Department of Neurology, Mayo Clinic Rochester, Minnesota USA; Department of Laboratory Medicine and Pathology, Mayo Clinic Jacksonville, Florida USA

**Keywords:** NULISA, proteomics, Alzheimer’s disease, young-onset, postmortem brain, tau, heterogeneity

## Abstract

**Background:** Alzheimer’s disease (AD) is clinicopathologically heterogeneous. A proportion of patients living with AD present clinically at a younger onset of cognitive symptoms before 65 years old and/or non-amnestic clinical syndromes. Neuropathologically, corticolimbic distribution of neurofibrillary tangle pathology occurs on a continuum with some cases having greater cortical tau pathology relative to limbic regions and others with relatively restricted accumulation in limbic structures. These patterns of corticolimbic tangle distribution are associated with clinical presentation and age at onset. This study sought to examine protein expression differences across the spectrum of clinicopathologic heterogeneity using the NULISA targeted proteomics platform.

**Methods:** A series of thirteen neuropathologically diagnosed AD cases from Mayo Clinic prospectively followed research studies were selected to reflect heterogeneity of clinical syndromes and corticolimbic distribution of tangle pathology. Frozen postmortem brain tissue samples were isolated from inferior parietal cortex and homogenized in RIPA buffer for analysis using Alamar Biosciences’ NULISA CNS disease 120 panel. Applying a conservative detection threshold of 75% level of detection for the novel application of NULISA in human brain, we evaluated levels of 69 of 129 protein targets across samples. We examined associations between age at onset cognitive symptoms and corticolimbic distribution of tangles (CLix) separately with individual protein targets using linear regression analysis.

**Results:** AD cases with a younger age at onset had higher measured levels of ubiquitin, while older age was associated with higher levels of total tau, CRH, and NPTX2. Investigations of corticolimbic heterogeneity revealed AD cases with lower CLix score (i.e., cortical predominant distribution of tau) had higher measured p-tau181, p-tau231, ubiquitin, and p62. AD cases with higher CLix (i.e., relative cortical sparing) had higher levels of total tau, CRH, NPTX2, MDH1, and HBA1. Brain-derived total tau consistently showed a stronger association in both models.

**Conclusion:** This work demonstrates the utility of postmortem proteomics for investigating biomarkers associated with AD clinicopathologic heterogeneity. We observed proteomic differences in synapse integrity, tau post-translational modification, and ubiquitination associated with age at symptomatic onset and corticolimbic distribution of tangle pathology.

## Introduction

Recent advances in plasma biomarker sensitivity in Alzheimer’s disease (AD) have enabled more precise measures of amyloid-β and tau associated with neuropathologic changes [1–3], informing diagnostics and clinical trial risk stratification [4–6]. Targeted proteomics platforms, like NUcleic acid Linked Immuno-Sandwich Assay (NULISA) sequencing technology are reported to enable high-sensitivity, low-volume analysis of plasma and other biofluids using an expanded protein panel [7–12]. Studies examining targeted biomarker differences between cognitively normal controls and individuals with mild cognitive impairment and dementia attributed to AD have identified distinct changes in protein markers of synaptic function, neurodegeneration, and neuroinflammation along the progression of AD pathogenesis [8, 12]. Although these studies provide critical insight into molecular alterations indirectly reflecting brain changes associated with distinct cognitive trajectories, there remains a need for postmortem validation in neuropathologically diagnosed AD cases. We reasoned that the NULISA technology was adaptable to brain tissue homogenate and that protein expression differences would emerge in association with clinicopathologic variables.

AD is heterogeneous in clinical presentation and neuropathology, expanding past the historical definition of a memory predominant syndrome in the presence of amyloid-β plaques and neurofibrillary tangle pathology to encompass a broader understanding of complexity [13]. The age at onset of sporadic AD can range dramatically, however the majority of cases present with an onset of cognitive symptoms after age 65 [14]. Cases with a before age 65 are more likely to have a non-amnestic clinical presentation, greater neuropathologic burden, and different neuroinflammatory profile compared to cases with a typical onset after 65 years [13, 15].

The effort to understand heterogeneity in AD is approached through multiple disciplines. Studies investigating neuropathologic heterogeneity have described a scaled continuum of neurofibrillary tangle pathology reflecting corticolimbic distribution of tangle pathology burden in the cortex relative to the hippocampus—corticolimbic index (CLix) [16–18]. Distinct morphologic differences in extracellular amyloid-β deposits were reported in atypical AD in addition to histopathologic differences in glial reactivity [17–19]. In the effort to better understand biological heterogeneity across clinicopathologic forms of AD we performed targeted fluid biomarker analysis to postmortem brain homogenates to elucidate biomarker, inflammatory, and molecular differences not observable through histopathologic characterization. We hypothesized that protein target variability measured in brain would associate with clinical measures (i.e., age at onset of cognitive symptoms) and neuropathologic heterogeneity (i.e. corticolimbic tangle distribution).

In the current era of disease-modifying therapeutics, there is an urgent need to better understand biological drivers of clinicopathologic heterogeneity in AD. In the present study, we sought to examine the protein level differences across different clinicopathologic forms of AD. In a pilot series of 13 neuropathologically characterized AD cases, we performed NULISA to examine protein-level differences in the AD brain and identified proteomic differences associated with age at symptomatic onset and neuropathologic distribution of neurofibrillary tangle pathology.

## Methods

### Defining clinicopathologic heterogeneity

We investigate clinical heterogeneity by examining age onset of cognitive symptoms, which range in our series from 51 to 83 years (Table 1). We examine age at onset and neuropathologic heterogeneity on a continuum. Neuropathologic heterogeneity was defined using the corticolimbic index (CLix) to spatially collapse corticolimbic tangle distribution into a score of 0-40, as previously described [17, 18]. The method relies upon thioflavin-S positive tangles quantified in three regions of the association cortices (middle frontal, superior temporal, and inferior parietal cortices) and two hippocampal subfields (CA1 and subiculum). Quantified tangle counts were normalized to the median tangle count in a large, postmortem AD series resulting in a normalized ratio of hippocampal to cortical tangle burden, ranging from 0 to 40—reflecting the continuum of corticolimbic tangle distribution.

**Table 1.** Case Characteristics.

| Case Number | Demographics and Clinical |  |  |  |  |  | Neuropathology |  |  |  |
| --- | --- | --- | --- | --- | --- | --- | --- | --- | --- | --- |
|  | Female Sex | APOE ɛ4 carriers | Age Onset, years | Age at death, years | Atypical | Clinical Syndrome | Braak stage | Thal phase | Brain weight (g) | CLix score |
|  | 5/13 (38%) | 10/13 (77%) | 63 (56,74) | 74 (62,82) | 6/13 (46%) |  | VI (V, VI) | 5 (4.5,5) | 1200 (1127,1328) | 13 (6.7,25) |
| 1 | F | 44 | 51 | 60 | Atypical | CBS | VI | 4 | 904 | 2.9 |
| 2 | M | 34 | 52 | 60 | Atypical | CBS | VI | 5 | 1148 | 7.2 |
| 3 | M | 33 | 53 | 59 | Amnestic | AD dementia | VI | 5 | 1291 | 6.7 |
| 4 | M | 44 | 59 | 66 | Atypical | PPA | VI | 5 | 1360 | 5.5 |
| 5 | M | 33 | 59 | 64 | Atypical | CBS | VI | 5 | NA | 3.4 |
| 6 | F | 44 | 62 | 75 | Amnestic | AD dementia | V | 5 | 1120 | 36 |
| 7 | M | 34 | 63 | 70 | Amnestic | AD dementia | VI | 5 | 1200 | 7.3 |
| 8 | F | 44 | 68 | 74 | Atypical | PCA | VI | 5 | 1340 | 15 |
| 9 | M | 34 | 69 | 77 | Amnestic | AD dementia | VI | 5 | 1070 | 32 |
| 10 | M | 34 | 73 | 78 | Atypical | CBS | V | 5 | 1200 | 13 |
| 11 | M | 34 | 75 | 85 | Amnestic | AD dementia | V | 5 | 1486 | 25 |
| 12 | F | 34 | 82 | 88 | Amnestic | AD dementia | VI | 3 | 1200 | 37 |
| 13 | F | 33 | 86 | 90+ | Amnestic | AD dementia | V | 4 | 1268 | 25 |
| Note: Data of continuous variables reported as median (interquartile range) for all neuropathologically diagnosed AD cases studied. Atypical clinical presentations included CBS, PCA, PPA. Braak tau staging reported in roman numerals. Abbreviations: AD= Alzheimer's disease dementia, CBS= corticobasal syndrome, CLix= corticolimbic index, F= female, g= gram, M= male, NA=not available, PCA= posterior cortical atrophy, PPA= primary progressive aphasia. |  |  |  |  |  |  |  |  |  |  |

### Case characteristics

Cases were enrolled in the Mayo Clinic Study of Aging (Rochester, MN) or the Mayo Clinic Alzheimer’s Disease Research Center (ADRC) longitudinal research study (Rochester, MN and Jacksonville, FL). We identified 38 cases with antemortem plasma collected within 3 years of death, CLix, frozen tissue, and documentation of clinical onset of cognitive symptoms. The broader project was designed for paired brain-plasma analyses and therefore cases with antemortem plasma proximate to death were prioritized. We selected 13 cases for this analysis based on range of age at onset (51-83 years) and CLix distribution (2.9-37) [17]. Eight cases were excluded because of missing documentation of age at the onset of cognitive symptoms. Written informed consent was obtained from all next of kin for participation in clinical research and brain autopsy. Ethical approval for this study was granted by the Mayo Clinic Institutional Review Board (IRB# 25-000355).

### Postmortem tissue preparation

Frozen tissue was sectioned from whole-hemisphere frozen brain tissue. The inferior parietal cortex was isolated based on identification of the angular gyrus. Gray matter was separated from white matter on dry ice to preserve protein and RNA integrity. Tissue integrity was assessed by RNA integrity number (RIN) with RIN above 6.0 measured for all samples included in analysis. A total of 500 mg of frozen tissue was collected from each sample. Frozen brain samples were homogenized in RIPA buffer (Pierce VWR, P189900) with protease inhibitor cocktail (Sigma-Aldrich, Complete Mini EDTA free 11836170001 and 1:100 phosphatase inhibitor cocktail 2 and 3 (Sigma-Aldrich, P5726-1mL and P0044-1mL) and centrifuged at 17,000 × g for 15 minutes. The supernatant of each sample was collected for proteomic analysis and aliquoted into 100µl aliquots.

### Sample analysis

RIPA prepared brain homogenate samples were run on the Alamar Bioscience NULISA CNS Disease Panel following published protocols [12]. In brief, RIPA soluble fraction isolated from frozen postmortem brain homogenates (n =13) were centrifuged at 17,000 × g for 15 minutes and the supernatant was extracted for analysis. Supernatant samples were normalized to a concentration of 1mg/mL and further diluted to a concentration of 0.02mg/mL with NULISA SD buffer. Supernatant samples and lysis buffer controls were incubated with a cocktail of capture and detection antibodies labeled with DNA barcodes. Immunocomplexes were purified, and cDNA sequences were generated by ligating DNA barcodes from antibody pairs. DNA levels were quantified by next-generation sequencing. Samples were run with two replicates of a plasma sample control, three inter-plate plasma controls, and four negative controls (sample buffer) to assess assay performance. Brain homogenate samples, plasma sample controls, inter-plate plasma controls, and buffer controls were processed on the same plate. Protein measurements were reported in NULISA protein quantification (NPQ) units. NPQ values were derived by log2 transformed raw values normalized to internal control measurements and divided by the median inter-plate control measurements.

### Neuropathologic assessment

Cases underwent standardized neuropathologic evaluation by a board-certified neuropathologist as previously described [20]. Upon brain donation, the left hemisphere was frozen in toto and stored at −80 °C for future biochemical studies; the right hemisphere was immediately processed for histologic evaluation. After two weeks fixation, tissue samples were collected according to the Consortium to Establish a Registry for Alzheimer’s Disease (CERAD) protocol [21]. Tissue blocks were embedded in paraffin and sectioned at a thickness of 5 µm, then mounted on positively charged microscopic slides for histologic and immunohistochemical evaluation. AD neuropathologic severity was assessed by Braak tau staging and amyloid-β by Thal phase [22, 23]

### Statistical Analysis

Data were analyzed using R (version 4.1.1). Data organization and statistical analyses were performed using R packages: readxl, dplyr, stats, labelled. Forest plot figures were created with ggplot2. Biomarker proteins measured in NPQ units are summarized as mean (standard deviation) and median (25th percentile, 75th percentile). Prior to modeling, biomarker proteins were standardized to have a mean of 0 and a standard deviation of 1. Univariate linear regression models were fitted for each biomarker protein and outcome (i.e., age at onset, CLix) to estimate the change in the outcome associated with a one–standard deviation increase in the biomarker protein. Linear regression models were used to evaluate associations between protein expression and continuous variables, i.e., age at onset and CLix. All statistical tests were two-sided, and p-values < 0.05 were considered statistically significant. * < 0.05, ** < 0.01, *** < 0.001. Nominal p-values were used to identify candidate protein associations.

## Results

Targeted proteomic analyses were performed on inferior parietal cortex homogenates from 13 sporadic autopsy confirmed AD cases to investigate NULISA application to postmortem brain, and association of individual protein targets with clinicopathologic variables of age of symptomatic onset (median=63 years [interquartile range 56-74]) and corticolimbic tangle distribution using CLix (13 [6.7-25]). Case characteristics are summarized in Table 1.

### NULISA readily detected targeted proteins in postmortem human brain homogenates

Table 2 summarizes target detection information. According to the manufacturer, protein targets were classified into the following groups: amyloid & tau, neurodegeneration, neuroinflammation, synuclein & synaptic proteins, and vascular & metabolism markers (Supplemental table S1).The CNS Disease panel includes 131 total protein targets, 129 detectable in brain. We detected 80/129 (62%) protein targets at or above a level of detection (LOD) of 50%. We opted to apply a conservative threshold of 75% LOD for the novel application of NULISA to brain homogenate, retaining proteins in our analysis if values were above the assay detection threshold in at least 75% of brain homogenate samples. Applying this conservative threshold detected 69/129 (53%) protein targets across all samples. Of the 69 protein targets we detected 13/22 (59%) amyloid & tau targets, 14/21 (67%) synuclein & synaptic proteins, 3/12 (25%) vascular & metabolism markers, 22/24 (92%) neurodegeneration markers, and 17/52 (33%) neuroinflammatory markers in postmortem brain samples. Notably, Aβ38, Aβ42, BDNF, and pTDP43-409 were detected at levels below the conservative 75% LOD threshold applied for this analysis, Figure 1. Given the low detectability of markers within the vascular & metabolism and neuroinflammatory categories, we restricted subsequent analyses to markers of: amyloid & tau pathology, synuclein & synaptic integrity, and neurodegeneration. The median coefficient of variation across sample controls was 4.99% (interquartile range [IQR], 3.3-7.4%).

**Figure 1.**
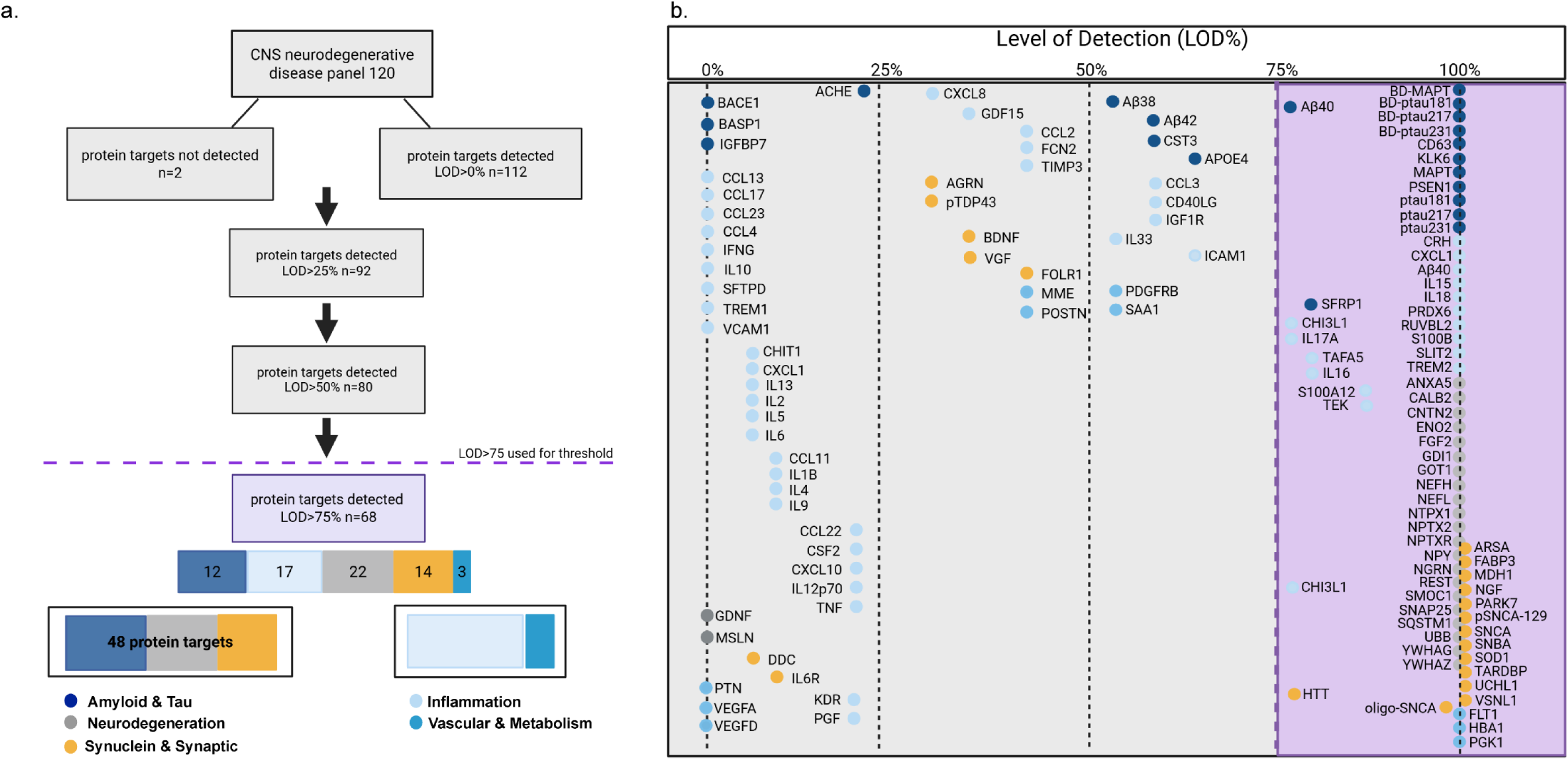
Workflow and detection of protein targets in postmortem brain homogenate. A) Workflow of protein target inclusion and exclusion. Β) Detection of individual protein targets.

**Table 2.**
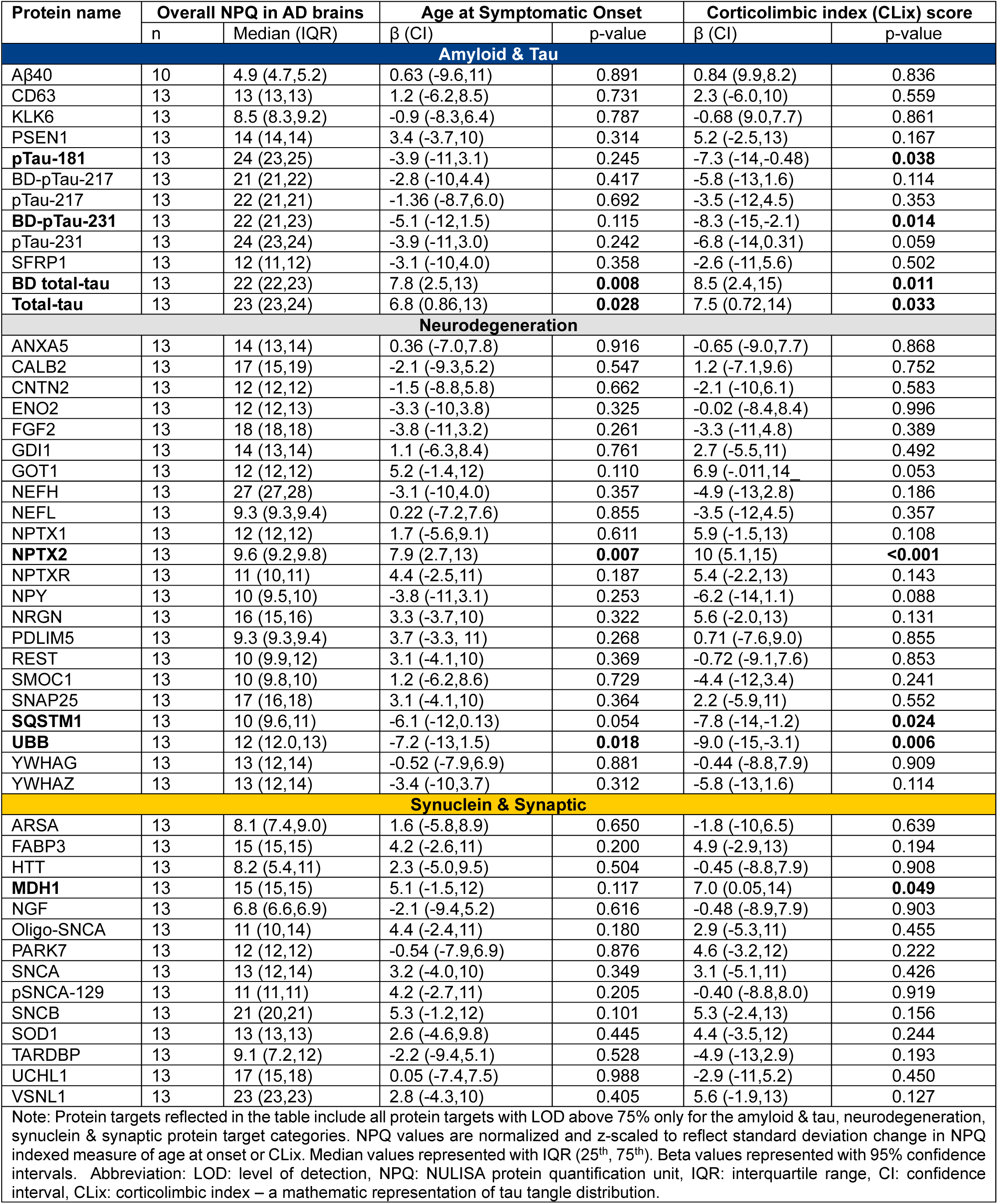

### Clinical heterogeneity in age at symptomatic onset shows significant association with ubiquitin, total tau, CRH, and NPTX2 protein levels

Protein expression of each target was examined in relation to age at symptomatic, Table 2. We observed differences in total tau (t-tau), a marker associated generally with synaptic degeneration [24, 25], and age at onset. Each standard deviation NPQ increase in t-tau was associated with a 6.8-year later age at onset (β = 6.8 [CI: 0.86, 13], p = 0.028). The brain-derived (BD) specific t-tau target demonstrated a greater effect with each standard deviation NPQ increase in BD t-tau was associated with a 7.8-year later onset of cognitive symptoms (β = 7.8 [CI: 2.5, 13], p = 0.008). Additionally, we observed associations between NPTX2 and ubiquitin with age at onset, Figure 2. We observed higher levels of NPTX2, a secreted neuronal protein involved in microglial-mediated synapse pruning [26], in cases with older symptomatic onset. Each standard deviation NPQ increase in NPTX2 was associated with 7.9-years older age at onset (β = 7.9 [CI: 2.7, 13], p = 0.007). Ubiquitin, a cellular marker for proteosome mediated degradation, was inversely associated with age at onset. Each standard deviation increase in ubiquitin was associated with an age at onset 7.2-years younger age at onset (β = -7.2 [CI: -13, -1.5], p = 0.018). Expression of corticotropin-releasing hormone (CRH) was positively associated with age at symptomatic onset; Supplemental Table S2, Supplemental Figure S1. Each standard deviation NPQ increase in CRH was associated with a 6.3-year later age at onset (β = 6.3 [CI: 0.18,12], p = 0.045). Confidence intervals were wide, likely due to the small sample size.

**Figure 2.**
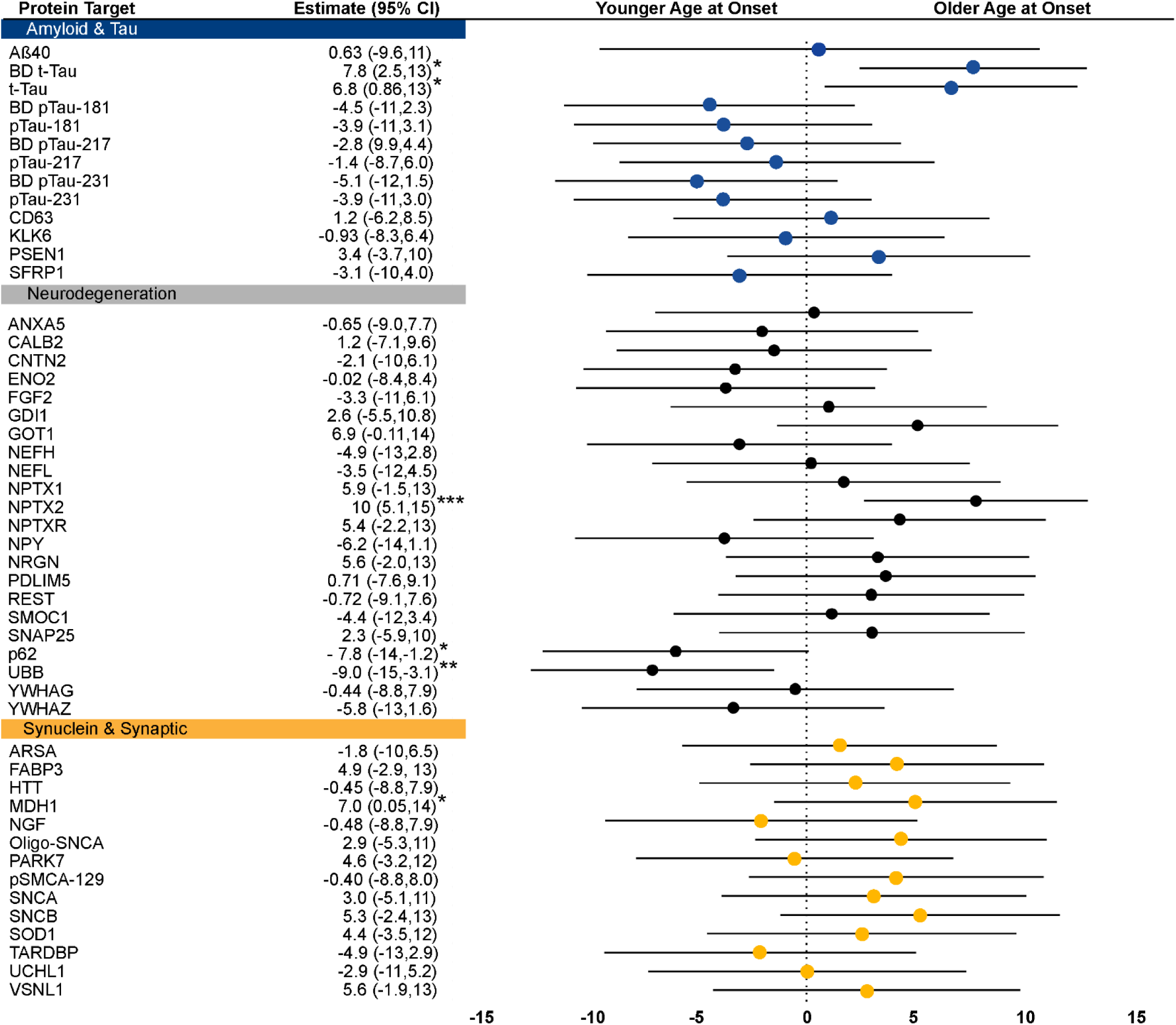
Forest plot of significantly associated protein targets and age onset. Association of BD-tau, tau, and NPTX2 with age at onset. β values derived from linear regression model of NPQ measured value and age at onset. Abbreviations: BD= brain derived, tau= microtubule associated protein tau (MAPT), NPTX2= neuropentraxin-2. Note: only associations between protein targets within amyloid & tau, neurodegeneration, and synuclein & synaptic categories were assessed.

### Neuropathologic heterogeneity in corticolimbic tangle distribution associated with tau posttranslational modifications, ubiquitin, p62, CRH, NPTX2, MDH1, and HBA1 protein levels

Clinical heterogeneity is paralleled microscopically by divergent tau patterns of neuropathologic tangle accumulation. We next analyzed proteomic differences along the continuum of neuropathologic heterogeneity using CLix to quantitatively assess corticolimbic tangle distribution [17]. Linear regression analysis revealed that each standard deviation NPQ increase in t-tau was associated with a 7.5-point increase in CLix (β = 7.5 [CI: 0.72, 14], p = 0.033). BD t-tau was associated with an 8.5-point higher CLix score (β = 8.5 [CI: 2.4, 15], p = 0.011). In contrast, each standard deviation NPQ increase in BD p-tau231 was associated with an 8.3-point lower CLix score (β = -8.3 [CI: -15, -2.0], p = 0.014). Each standard deviation NPQ increase in BD pTau181 was associated with 7.9-points lower CLix score (β = -7.9 [CI: -14, -1.3], p = 0.022), with non-BD pTau181 similarly associated with a lower CLix score (β = -7.3 [CI: -14, -0.48], p = 0.038); Table 2, Figure 3.

**Figure 3.**
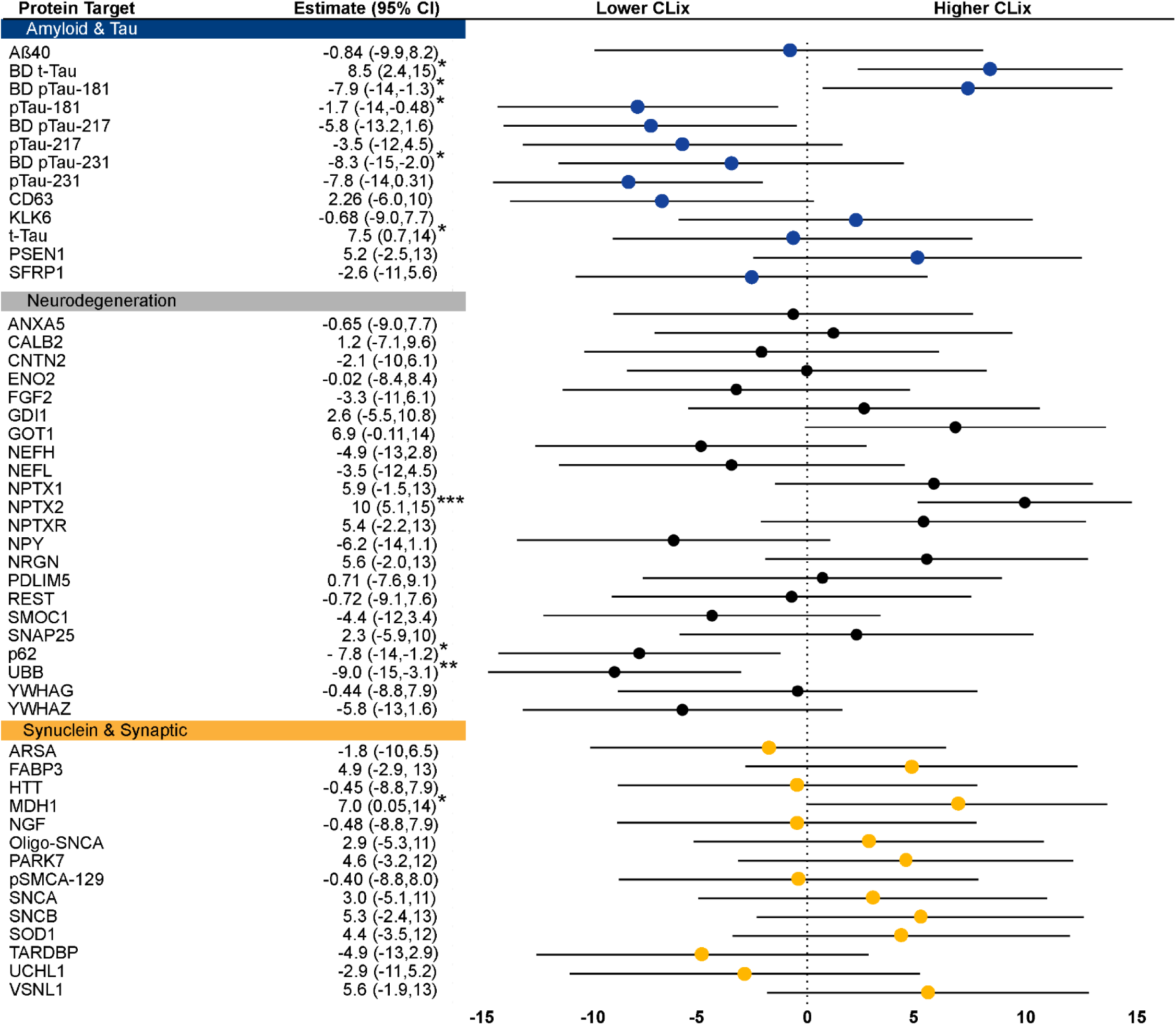
Forest plot of significantly associated protein targets and CLix Forest plot of significantly associated protein targets and corticolimbic tangle distribution. β values derived from linear regression model of NPQ measured value and CLix. Abbreviations: BD= brain derived, tau= microtubule associated protein tau (MAPT), MDH1=malate dehydrogenase-1, NPTX2= neuropentraxin-2, UBB= ubiquitin. Note: only associations between protein targets within amyloid & tau, neurodegeneration, and synuclein & synaptic categories were assessed.

We observed associations between three proteins and corticolimbic tangle distribution: p62, ubiquitin, and NPTX2 (Figure 3). In cases with lower CLix scores or those with greater cortical tangle distribution, we observed higher p62 and ubiquitin protein levels. Each standard deviation NPQ increase in p62 was associated with a 7.8-points lower CLix score (β = -7.8 [CI: -14, -1.2], p = 0.024). Each standard deviation NPQ increase in ubiquitin was associated with an 9.0-point lower CLix score (β = -9.0 [CI: -15, -3.1], p = 0.006). Conversely, we observed higher expression of NPTX2 in cases with higher CLix scores. Each standard deviation NPQ increase in NPTX2 was associated with a 10.1-point higher CLix score (β = 10 [CI: 5.1, 15], p < 0.001).

Additionally, we observed associations between CLix and malate dehydrogenase-1 (MDH1), a cytosolic enzyme integral to mitochondrial NADH supply and oxidative phosphorylation (ox-phos) and hemoglobin subunit α-1 (HBA1), a protein involved with oxygen transport [27–29]. MDH1 was higher in cases with higher CLix scores. Each standard deviation increase in MDH1 was associated with a 7.0-point higher CLix score (β = 7.0 [CI: 0.050, 14], p = 0.029). HBA1 expression was higher in cases with higher CLix scores with each standard deviation NPQ increase in HBA1 associated with 9.1-points higher CLix (β=9.1 [CI:3.3,15], p=0.005), Supplemental Figure S2.

## Discussion

In this pilot postmortem AD series, targeted multiplex proteomic profiling of inferior parietal cortex identified protein-level differences associated with age at onset and corticolimbic tau distribution, supporting further consideration of multiplex proteomics to study biological heterogeneity in AD. AD heterogeneity is reflected during life with different clinical symptoms and trajectories and heterogeneity is also reflected neuropathologically [13, 30]. We reasoned that fluid-based biomarker technology could be applied to postmortem human AD brains to investigate protein-level differences associated with measurable clinicopathologic heterogeneity.

Applying a conservative detectability cutoff of 75%, we detected 53% of protein targets in postmortem brain homogenate. Examining the association between individual protein targets and clinicopathologic variables, we found younger age onset and lower CLix (i.e., cortical predominant tangle distribution) associated with lower total tau, lower NPTX2, and higher ubiquitin protein levels. Lower CLix was additionally associated with higher phospho-tau (BDpTau-181, pTau-181, BD-pTau-231) and lower levels of the ox-phos regulating protein MDH1. Our findings provide supportive evidence for biologic differences relevant to AD clinicopathologic heterogeneity.

Neuropathologic diagnostics are often limited to routine diagnostic immunohistochemistry. To explore the utility of fluid-based biomarker technology for neuropathologic analysis of postmortem human brain, we applied NULISA to parietal cortex homogenates to measure protein-level differences across clinicopathologic forms of AD. NULISA requires a low sample volume, thereby maximizing the amount of analysis that can be performed per sample. We were able to assess several AD relevant biomarkers commonly used in the AD biomarker field to clinically evaluate, detect, and monitor AD in vivo including t-tau, p-tau 181, p-tau 217, p-tau 231, GFAP, NfL, and YKL-40 [7]. All were detected in our postmortem brain homogenates. The low detectability of neuroinflammatory markers in our samples suggests that the ability to detect differences in inflammatory markers using RIPA digested postmortem brain homogenate may be limited in the current protocol and should be further investigated to improve detection (e.g., lysis buffer, spin duration, homogenate concentration).

In the present study we examined protein expression in relation to age onset of cognitive symptoms, corticolimbic distribution of tangle pathology (CLix). Age onset, used as a continuous variable, allowed us to examine cases on a continuum rather than applying a nominal boundary of 65 years to distinguish “young-onset” and “late-onset” presentations. We observed associations between age at onset and t-tau, NPTX2, and ubiquitin. BD t-tau, t-tau, NPTX2, and were higher in cases with older age at onset. Higher levels of t-tau in plasma and CSF of healthy older adults and AD participants suggest an age associated increase in t-tau independent of the AD pathologic cascade [31, 32]. In the present study we observed a positive correlation between age at onset and NPTX2 protein levels. A recent examination of plasma study identified 17 protein targets associated with age, not including NPTX2 [12]. CSF proteomic studies have shown an association between NPTX2 and cortical thickness [33, 34]. Moreover, in a study using both animal and cell culture models found that overexpression of NPTX2 was sufficient to reduce complement activity and mitigate synapse loss together providing promising evidence to support that levels of NPTX2 may be indicative of neuronal integrity or brain health [26]. Consistently, biofluid studies have reported lower NPTX2 in AD [35, 36] however this is the first study of which we are aware that has described intra-disease differences in NPTX2 levels associated with clinicopathologic phenotypes of AD.

We observed associations with clinicopathologic variables and ubiquitin. Ubiquitin was higher in cases with a younger age at onset. Collectively, these findings suggest that younger onset AD may have greater ubiquitin-proteosome dysfunction compared to cases with later onset. Dysregulated ubiquitin proteosome system (UPS) is a well-established phenomenon in AD occurring early in the AD process [37] however differences in UPS among clinical variants in AD are underexplored. Our findings of greater ubiquitin with unchanged UCHL1 is suggestive of greater protein ubiquitination or alternatively reduced degradation (i.e. greater UPS dysfunction) specifically in YOAD. These findings are also consistent with neuropathologic and imaging studies describing greater neurodegeneration and neuropathologic burden in YOAD [38–40].

Approaching AD heterogeneity through the lens of neuropathology, we examined protein-level differences along the CLix continuum, a measure of corticolimbic tangle distribution [17]. There is significant overlap between cases with atypical neuropathologic accumulation of tau (i.e., cortical predominant/hippocampal sparing and limbic predominant) and younger age at onset of cognitive symptoms. This has been reported previously and was purposefully reflected in the heterogeneity in the present series [16–18, 41]. AD cases within the present series ranged in CLix from 2.9 to 37, we observed an association between CLix and p-tau with greater BD p-tau231 and p-tau181 observed in cases with lower CLix score or greater cortical tangle pathology. These findings expand on the definition of AD neuropathologic subtypes, that are derived from Thioflavin-S tangle counts to characterize the specific p-tau isoforms and biological differences associated with different neuropathologic distributions.

Biofluid and neuroimaging studies generally regard p-tau181 and p-tau231 as reflective of AD neuropathology [42, 43]. These fluid biomarkers are less correlated with tau PET uptake and more ascribed to different stages of tangle maturity: postmortem histological validation studies show p-tau181 is present in pre-tangles and p-tau231 in mature tangles, neuritic lesions, and described in the axonal compartment [44, 45]. Colocalization studies have localized p-tau231 to synapses and observed overlap between p-tau231 and PSD-95, a marker of postsynaptic density of excitatory synapses [46]. In the context of our findings across clinical syndromes, age at onset, and neuropathologic heterogeneity, the observed differences in p-tau181 and p-tau231 suggest possible differences in neuritic pathology and synaptic integrity across clinicopathologic forms of AD.

Ubiquitin was also observed to associate with corticolimbic tangle distribution. Specifically, we observed a strong inverse association with ubiquitin levels and CLix. Dysregulation of the ubiquitin-proteosome system (UPS) and ubiquitination in AD is well documented [29, 47]. Further, ubiquitin is a common posttranslational modification of tau, playing a role in tau degradation and aggregation of insoluble p-tau [48, 49]. However, the biological consequence of ubiquitin is largely context dependent as ubiquitin is a molecular signal common to both the UPS and autophagy pathways [50]. Here we observed greater ubiquitin in cases with lower CLix, a pattern congruent with what we observed with measured p-tau181 and p-tau231. Taken together these results are consistent with histopathologic evaluations of tangle maturity along the CLix continuum with greater advanced tangle pathology reported in the inferior parietal cortex specifically in cases with lower CLix compared to cases with highest CLix scores [17]. Further examination of UPS markers and markers of autophagy, comparing burden and activation across multiple brain regions, are necessary to probe the mechanistic link between these two observations in cases with cortical predominant tangle distribution.

Moreover, we observed greater HBA1, NPTX2, and MDH1 in cases with higher CLix scores, suggesting that regions with less tangle accumulation (i.e., the inferior parietal cortex of these cases) may have less oxidative stress, and synaptic dysfunction as the brain homogenates assessed in this study were sampled from the inferior parietal cortex [51, 52]. There is limited evidence to interpret the HBA1 finding within the context of AD however to date several studies have reported higher HBA1 protein expression in plasma of probable AD cases [10, 53]. Multiple human postmortem proteomics studies have also reported higher HBA1 expression in AD cases compared to controls [29, 54, 55]. Drummond et al did not observe interaction between HBA1 and tangles but reported HBA1 in laser capture microdissected neurofibrillary tangles [56]. Ingenuity protein analysis revealed tau as an upstream regulator of HBA1 [56]. Evidence from animal studies suggests that NPTX2 regulates microglial activation and microglia mediated synapse loss [26]. Human postmortem transcriptomic studies reflect reduced expression of NPTX2 in the AD brain across frontal, temporal, and parietal cortices. NPTX2 levels in CSF have been associated with resistance to brain atrophy and cognitive resilience in longitudinal studies evaluating brain atrophy rates of cognitively unimpaired longitudinally followed participants [33, 34]. While all cases in our series had cognitive impairment and advanced AD neuropathologic change these differences in NPTX2 suggest possible regional differences in neuronal health and synaptic in integrity in these cases. Levels of MDH1 are reported to be lower in AD compared to healthy controls [28]. Here in our intra-disease analyses we observed greater MDH1 levels in cases with higher CLix scores (i.e., limbic predominant tangle distribution and less cortical tangle pathology) suggesting a potentially greater level of ox-phos in regions of greater tangle pathology [12, 28].

Limitations of this study include the small case series as this limits the generalizability of these findings and underscores the need for replication of these methods in a larger series using non-overlapping comparative groups. This study was designed to investigate intra-disease protein differences within the disease context of AD heterogeneity and therefore did not include non-AD controls. Future studies would benefit from the inclusion of non-AD neurodegenerative and non-diseased controls to better evaluate proteomic differences within and outside of the context of neurodegeneration. Nominal p-values were used to identify candidate protein associations with continuous variables of age at onset and corticolimbic heterogeneity but should not be interpreted as confirmatory as orthogonal validation of candidate proteins are needed. Expanding the sample and addition of comparative groups would allow for further examination of plausible neuroprotective markers, including NPTX2, CRH, and HBA1, outside the context of disease, where age-associated changes remain poorly understood. Lastly, the present study was performed in a single brain region, utilizing plasma samples as internal plate controls, limiting interpretation of regional differences in protein expression across the brain. The analysis was performed on the RIPA-soluble fraction of inferior parietal cortex, measured protein levels may reflect soluble protein abundance, extraction efficiency, and epitope availability rather than total tissue burden. Future studies extending this analysis of clinicopathologic heterogeneity in AD to plasma are planned to explore potential biomarkers capable of differentiating clinicopathologic subtypes of AD during life.

## Abbreviations

Aβ: Amyloid-beta
AD: Alzheimer’s disease
ADRC: Alzheimer’s Disease Research Center
APOE: Apolipoprotein E
BD: Brain derived
BDNF: Brain-derived neurotrophic factor
CBS: Corticobasal syndrome
CERAD: Consortium to Establish a Registry for Alzheimer’s Disease
CHI3L1: Chitinase-3-like protein 1 aka YKL-40
CLix: Corticolimbic index
CNS: Central nervous system
CRH: Corticotropin-releasing hormone
CSF: Cerebrospinal fluid
HBA1: Hemoglobin subunit alpha 1
IPA: Ingenuity protein analysis
LOAD: Late-onset Alzheimer’s disease
MDH1: Malate dehydrogenase 1
NGS: Next-generation sequencing
NPQ: NULISA protein quantification
NPTX2: Neuronal pentraxin-2
NULISA: nucleic acid linked immuno-sandwich assay
Ox-phos: oxidative phosphorylation
PCA: Posterior cortical atrophy
PET: Positron emission tomography
PPA: Primary progressive aphasia
p-tau: Phosphorylated tau
RIPA: radioimmunoprecipitation assay buffer
RNA: Ribonucleic acid
UCHL1: Ubiquitin C-terminal hydrolase L1
UPS: Ubiquitin-proteasome system
YOAD: Young-onset Alzheimer’s disease

## Declarations

This study was approved by the Ethical Review Board (IRB# 25-000355) at Mayo Clinic and conducted in accordance with the Declaration of Helsinki. Written informed consent was obtained from participants or next of kin, and appropriate authorization was granted through the Mayo Clinic Research Executive Committee for all research performed on *postmortem* brain samples. Requests for raw or processed from qualified investigators will undergo review by Mayo Clinic’s Legal Department to determine if confidentiality restrictions apply. Any data deemed eligible for sharing will be made available through a Data Use Agreement.

## Acknowledgements

We are grateful for the brain donors and their families and their commitment to research and scientific discovery. Our sincere thanks to the Alamar Biosciences Technology Access Program (TAP) for generously processing samples through the Technology Access Program. We note that Alamar Biosciences had no role in study design, statistical analysis, biological interpretation, manuscript drafting, or the decision to submit. We thank Ashley Wood and Monica Castanedes-Casey for their dedication and expertise. We are grateful for the attention to detail of our programmatic staff Sabrina Rothberg, Avery Hatfield, and Kelsey Caetano-Anolles.

## Competing interest, author contributions

Sara Rose Dunlop, PhD contributed to study conceptualization, study design, data acquisition, data analysis, data interpretation, manuscript writing, and manuscript editing. SRD has no competing interests.

Sarah J. Lincoln provided technical support and specimen processing. SJL has no competing interests.

Zhongwei Peng, MS performed statistical analyses. ZP has no competing interests.

Neill R. Graff-Radford, MBBCh, FRCP; Christian Lachner, MD; Gregory S. Day, MD; Ronald C. Petersen; Bradley

F. Boeve, MD; and Jonathan Graff-Radford, MD contributed to patient clinical evaluation and care.

NRGR has taken part in multicentre trials supported by Eli Lilly, Biogen, Eisai and Cognition Therapeutics which is outside the submitted work He has received publishing royalties from UpToDate, Inc for a chapter on NPH. CL has no competing interests. GSD reports no competing interests directly relevant to this work. His research is supported by NIH (R01AG089380, U01AG057195, U01NS120901, U19AG032438, P30AG062677). He serves as a Topic Editor (Dementia) for DynaMed (EBSCO). He is a co-Project PI for a clinical trial in anti-NMDAR encephalitis, which receives support from NIH/NINDS (U01NS120901) and Amgen Pharmaceuticals. He has developed educational materials for Continuing Education Inc, Ionis Pharmaceuticals, and MJH Life Sciences. He owns stock in ANI Pharmaceuticals. GSD (and Mayo Clinic) has received in-kind contributions for radiotracer precursors for tau-PET neuroimaging in studies of memory and aging (via Avid Radiopharmaceuticals, a wholly owned subsidiary of Eli Lilly).

RCP Dr. Ron Petersen serves as a consultant for Biogen, Inc., Roche, Inc., Merck, Inc., Genentech Inc. (DSMB), Nestle, Inc., Eli Lilly and Co., Novartis, Novo Nordisk and Eisai, Inc., receives publishing royalties from Mild Cognitive Impairment (Oxford University Press, 2003), and UpToDate.

BFB receives research support for clinical trials sponsored by Alector, Cognition Therapeutics, EIP Pharma/Cervomed, and Transposon. He serves on the Scientific Advisory Board of the Tau Consortium – funded by the Rainwater Charitable Foundation. JGR is a site investigator for clinical trials sponsored by Eisai and Cognition therapeutics, serves on the DSMB for StrokeNET, is an associate editor for JAMA Neurology and receives honoraria for serving as faculty at IMPACT AD and the American Academy of Neurology.

Jessica F. Tranovich provided editorial and programmatic support. JFT has no competing interests.

R. Ross Reichard MD, Dennis W. Dickson MD, Aivi T. Nguyen MD contributed to neuropathologic diagnosis of cases. RRR, ATN report no disclosures. DWD receives research support from the NIH (P30 -G062677; P50-NS072187; P01-AG003949) from the Mangurian Foundation Lewy Body Dementia Program at Mayo Clinic and the Robert E. Jacoby Professorship; is an editorial board member of Acta Neuropathologica, Annals of Neurology, Brain, Brain Pathology, and Neuropathology, and he is editor in chief of American Journal of Neurodegenerative Disease. Dr. Lea T. Grinberg received honorarium for education material from Orator Inc and University of California Berkeley. LTG is supported by grant from the CureAlz Fund, Ed and Ethyl Moore, Rainwater Charitable Foundation, and by the NIH/National Institute on Aging (R01-AG075802, R01-AG064314, R01-AG070826, R01-AG060477, U54NS123746, K24-AG053435. LTG serves on the MSAG for the Alzheimer’s Association and governing board of the Global Brain Health Institute.

Alicia Algeciras-Schimnich, PhD contributed expertise in clinical chemistry and biomarker interpretation.

AAS has participated in advisory boards for Roche Diagnostics, Fujirebio Diagnostics, Siemens Healthineers, and Beckman Coulter; and has received speaker honoraria from Roche Diagnostics, Beckman Coulter and Eli Lilly.

Melissa E. Murray, PhD served as principal investigator, contributed to funding acquisition, study conceptualization and design, and manuscript editing. MEM received grant funding from Eli Lilly and company and speaker fees from Eisai. MEM is supported by grants from the Alzheimer’s Association Florida Gulf Coast Chapter (SG-25-1416824), Rainwater Charitable Foundation, and by the NIH/National Institute on Aging (R01-AG075802, R01-AG073282, U01-AG057195, P30-AG062677, U19-AG069701, RF1 AG069052).

## References

1. Ashton NJ, Pascoal TA, Karikari TK, Benedet AL, Lantero-Rodriguez J, Brinkmalm G, Snellman A, Scholl M, Troakes C, Hye A, et al: Plasma p-tau231: a new biomarker for incipient Alzheimer’s disease pathology. Acta Neuropathol 2021, 141:709–724.

2. Bermudez C, Graff-Radford J, Syrjanen JA, Stricker NH, Algeciras-Schimnich A, Kouri N, Kremers WK, Petersen RC, Jack CR, Jr., Knopman DS, et al: Plasma biomarkers for prediction of Alzheimer’s disease neuropathologic change. Acta Neuropathol 2023, 146:13–29.

3. Hu M, Moloney CM, Przybelski SA, Algeciras-Schimnich A, Therneau TM, Fought AJ, Rothberg DM, Nguyen AT, Reichard RR, Dickson DW, et al: Association between antemortem plasma and structural MRI biomarkers and postmortem tau pathology in the Mayo Clinic Study of Aging. Alzheimers Res Ther 2025, 17:224.

4. Devanarayan V, Charil A, Horie K, Doherty T, Llano DA, Andreozzi E, Sachdev P, Ye Y, Murali LK, Zhou J, et al: Plasma pTau217 ratio predicts continuous regional brain tau accumulation in amyloid-positive early Alzheimer’s disease. Alzheimers Dement 2025, 21:e14411.

5. Palmqvist S, Tideman P, Mattsson-Carlgren N, Schindler SE, Smith R, Ossenkoppele R, Calling S, West T, Monane M, Verghese PB, et al: Blood Biomarkers to Detect Alzheimer Disease in Primary Care and Secondary Care. JAMA 2024, 332:1245–1257.

6. Piura YD, Figdore DJ, Lachner C, Bornhorst J, Algeciras-Schimnich A, Graff-Radford NR, Day GS: Diagnostic performance of plasma p-tau217 and Abeta42/40 biomarkers in the outpatient memory clinic. Alzheimers Dement 2025, 21:e70316.

7. Feng W, Beer JC, Hao Q, Ariyapala IS, Sahajan A, Komarov A, Cha K, Moua M, Qiu X, Xu X, et al: NULISA: a proteomic liquid biopsy platform with attomolar sensitivity and high multiplexing. Nat Commun 2023, 14:7238.

8. Di Molfetta G, Pola I, Tan K, Isaacson R, Blennow K, Ashton NJ, Benedet AL, Zetterberg H: Inflammation biomarkers and Alzheimer’s disease: A pilot study using NULISAseq. Alzheimers Dement (Amst) 2025, 17:e70079.

9. DuBois KN, Pal S, Maher AC, Heidebrink J, Persad C, Giordani BM, Hampstead BM, Bakulski KM, Morgan DG, Kanaan NM: Dementia etiology classification using NULISA plasma biomarkers and machine learning. medRxiv 2025.

10. Ibanez L, Liu M, Beric A, Timsina J, Kohlfeld P, Bergmann K, Lowery J, Sykora N, Sanchez-Montejo B, Brock W, et al: Benchmarking of a multi-biomarker low-volume panel for Alzheimer’s disease and related dementia research. Alzheimers Dement 2025, 21:e14413.

11. Rea Reyes RE, Wilson RE, Langhough RE, Studer RL, Jonaitis EM, Oomens JE, Planalp EM, Bendlin BB, Chin NA, Asthana S, et al: Targeted proteomic biomarker profiling using NULISA in a cohort enriched with risk for Alzheimer’s disease and related dementias. Alzheimers Dement 2025, 21:e70166.

12. Zeng X, Sehrawat A, Lafferty TK, Chen Y, Rawat M, Kamboh MI, Villemagne VL, Lopez OL, Cohen AD, Karikari TK: Novel plasma biomarkers of amyloid plaque pathology and cortical thickness: Evaluation of the NULISA targeted proteomic platform in an ethnically diverse cohort. Alzheimers Dement 2025, 21:e14535.

13. Graff-Radford J, Yong KXX, Apostolova LG, Bouwman FH, Carrillo M, Dickerson BC, Rabinovici GD, Schott JM, Jones DT, Murray ME: New insights into atypical Alzheimer’s disease in the era of biomarkers. Lancet Neurol 2021, 20:222–234.

14. Kvello-Alme M, Brathen G, White LR, Sando SB: The Prevalence and Subtypes of Young Onset Dementia in Central Norway: A Population-Based Study. J Alzheimers Dis 2019, 69:479–487.

15. Thapa S, Anastassiadis C, Vasilevskaya A, Taghdiri F, Jurisica I, Hadian M, Salwierz P, Robbani F, Kivisakk P, Hyman B, et al: Distinct inflammatory profiles in young-onset versus late-onset Alzheimer’s disease. Alzheimers Dement 2025, 21:e70509.

16. Janocko NJ, Brodersen KA, Soto-Ortolaza AI, Ross OA, Liesinger AM, Duara R, Graff-Radford NR, Dickson DW, Murray ME: Neuropathologically defined subtypes of Alzheimer’s disease differ significantly from neurofibrillary tangle-predominant dementia. Acta Neuropathol 2012, 124:681–692.

17. Kouri N, Frankenhauser I, Peng Z, Labuzan SA, Boon BDC, Moloney CM, Pottier C, Wickland DP, Caetano-Anolles K, Corriveau-Lecavalier N, et al: Clinicopathologic Heterogeneity and Glial Activation Patterns in Alzheimer Disease. JAMA Neurol 2024, 81:619–629.

18. Murray ME, Graff-Radford NR, Ross OA, Petersen RC, Duara R, Dickson DW: Neuropathologically defined subtypes of Alzheimer’s disease with distinct clinical characteristics: a retrospective study. Lancet Neurol 2011, 10:785–796.

19. Boon BDC, Hoozemans JJM, Lopuhaa B, Eigenhuis KN, Scheltens P, Kamphorst W, Rozemuller AJM, Bouwman FH: Neuroinflammation is increased in the parietal cortex of atypical Alzheimer’s disease. J Neuroinflammation 2018, 15:170.

20. Murray ME, Moloney CM, Kouri N, Syrjanen JA, Matchett BJ, Rothberg DM, Tranovich JF, Sirmans TNH, Wiste HJ, Boon BDC, et al: Global neuropathologic severity of Alzheimer’s disease and locus coeruleus vulnerability influences plasma phosphorylated tau levels. Mol Neurodegener 2022, 17:85.

21. Mirra SS, Heyman A, McKeel D, Sumi SM, Crain BJ, Brownlee LM, Vogel FS, Hughes JP, van Belle G, Berg L: The Consortium to Establish a Registry for Alzheimer’s Disease (CERAD). Part II. Standardization of the neuropathologic assessment of Alzheimer’s disease. Neurology 1991, 41:479–486.

22. Braak H, Braak E: Neuropathological stageing of Alzheimer-related changes. Acta Neuropathol 1991, 82:239–259.

23. Thal DR, Rub U, Orantes M, Braak H: Phases of A beta-deposition in the human brain and its relevance for the development of AD. Neurology 2002, 58:1791–1800.

24. Soares C, Bellaver B, Ferreira PCL, Povala G, Schaffer Aguzzoli C, Ferrari-Souza JP, Zalzale H, Lussier FZ, Rohden F, Abbas S, et al: CSF total tau as a proxy of synaptic degeneration. Nat Commun 2025, 16:8076.

25. Samgard K, Zetterberg H, Blennow K, Hansson O, Minthon L, Londos E: Cerebrospinal fluid total tau as a marker of Alzheimer’s disease intensity. Int J Geriatr Psychiatry 2010, 25:403–410.

26. Zhou J, Wade SD, Graykowski D, Xiao MF, Zhao B, Giannini LAA, Hanson JE, van Swieten JC, Sheng M, Worley PF, Dejanovic B: The neuronal pentraxin Nptx2 regulates complement activity and restrains microglia-mediated synapse loss in neurodegeneration. Sci Transl Med 2023, 15:eadf0141.

27. Broeks MH, Shamseldin HE, Alhashem A, Hashem M, Abdulwahab F, Alshedi T, Alobaid I, Zwartkruis F, Westland D, Fuchs S, et al: MDH1 deficiency is a metabolic disorder of the malate-aspartate shuttle associated with early onset severe encephalopathy. Hum Genet 2019, 138:1247–1257.

28. Jia D, Wang F, Yu H: Systemic alterations of tricarboxylic acid cycle enzymes in Alzheimer’s disease. Front Neurosci 2023, 17:1206688.

29. Cunningham A, Barrett E, Risch S, Lee PHU, Lee C, Moghekar A, Patra P, Shim JW: NFkappaB1: a common biomarker linking Alzheimer’s and Parkinson’s disease pathology. Front Neurosci 2025, 19:1589857.

30. Sirkis DW, Bonham LW, Johnson TP, La Joie R, Yokoyama JS: Dissecting the clinical heterogeneity of early-onset Alzheimer’s disease. Mol Psychiatry 2022, 27:2674–2688.

31. Chatterjee S, Sealey M, Ruiz E, Pegasiou CM, Brookes K, Green S, Crisford A, Duque-Vasquez M, Luckett E, Robertson R, et al: Age-related changes in tau and autophagy in human brain in the absence of neurodegeneration. PLoS One 2023, 18:e0262792.

32. Chiu MJ, Fan LY, Chen TF, Chen YF, Chieh JJ, Horng HE: Plasma Tau Levels in Cognitively Normal Middle-Aged and Older Adults. Front Aging Neurosci 2017, 9:51.

33. Soldan A, Oh S, Ryu T, Pettigrew C, Zhu Y, Moghekar A, Xiao MF, Pontone GM, Albert M, Na CH, Worley P: NPTX2 in Cerebrospinal Fluid Predicts the Progression From Normal Cognition to Mild Cognitive Impairment. Ann Neurol 2023, 94:620–631.

34. Vazquez JP, Pettigrew C, Zhu Y, Anderson C, Erus G, Davatzikos C, Miller M, Moghekar A, Oh S, Na CH, et al: CSF Levels of NPTX2 Are Associated With Less Brain Atrophy Over Time in Cognitively Unimpaired Individuals. Ann Clin Transl Neurol 2025.

35. Belbin O, Xiao MF, Xu D, Carmona-Iragui M, Pegueroles J, Benejam B, Videla L, Fernandez S, Barroeta I, Nunez-Llaves R, et al: Cerebrospinal fluid profile of NPTX2 supports role of Alzheimer’s disease-related inhibitory circuit dysfunction in adults with Down syndrome. Mol Neurodegener 2020, 15:46.

36. Xiao MF, Xu D, Craig MT, Pelkey KA, Chien CC, Shi Y, Zhang J, Resnick S, Pletnikova O, Salmon D, et al: NPTX2 and cognitive dysfunction in Alzheimer’s Disease. Elife 2017, 6.

37. Zhang Y, Chen X, Zhao Y, Ponnusamy M, Liu Y: The role of ubiquitin proteasomal system and autophagy-lysosome pathway in Alzheimer’s disease. Rev Neurosci 2017, 28:861–868.

38. Marshall GA, Fairbanks LA, Tekin S, Vinters HV, Cummings JL: Early-onset Alzheimer’s disease is associated with greater pathologic burden. J Geriatr Psychiatry Neurol 2007, 20:29–33.

39. Scholl M, Ossenkoppele R, Strandberg O, Palmqvist S, Swedish Bio Fs, Jogi J, Ohlsson T, Smith R, Hansson O: Distinct 18F-AV-1451 tau PET retention patterns in early- and late-onset Alzheimer’s disease. Brain 2017, 140:2286–2294.

40. Spina S, La Joie R, Petersen C, Nolan AL, Cuevas D, Cosme C, Hepker M, Hwang JH, Miller ZA, Huang EJ, et al: Comorbid neuropathological diagnoses in early versus late-onset Alzheimer’s disease. Brain 2021, 144:2186–2198.

41. Boon BDC, Labuzan SA, Peng Z, Matchett BJ, Kouri N, Hinkle KM, Lachner C, Ross OA, Ertekin-Taner N, Carter RE, et al: Retrospective Evaluation of Neuropathologic Proxies of the Minimal Atrophy Subtype Compared With Corticolimbic Alzheimer Disease Subtypes. Neurology 2023, 101:e1412–e1423.

42. Mila-Aloma M, Ashton NJ, Shekari M, Salvado G, Ortiz-Romero P, Montoliu-Gaya L, Benedet AL, Karikari TK, Lantero-Rodriguez J, Vanmechelen E, et al: Plasma p-tau231 and p-tau217 as state markers of amyloid-beta pathology in preclinical Alzheimer’s disease. Nat Med 2022, 28:1797–1801.

43. Gonzalez-Ortiz F, Kirsebom BE, Contador J, Tanley JE, Selnes P, Gisladottir B, Palhaugen L, Suhr Hemminghyth M, Jarholm J, Skogseth R, et al: Plasma brain-derived tau is an amyloid-associated neurodegeneration biomarker in Alzheimer’s disease. Nat Commun 2024, 15:2908.

44. Moloney CM, Labuzan SA, Crook JE, Siddiqui H, Castanedes-Casey M, Lachner C, Petersen RC, Duara R, Graff-Radford NR, Dickson DW, et al: Phosphorylated tau sites that are elevated in Alzheimer’s disease fluid biomarkers are visualized in early neurofibrillary tangle maturity levels in the post mortem brain. Alzheimers Dement 2023, 19:1029–1040.

45. Wennstrom M, Schultz N, Gallardo PM, The Netherlands Brain B, Serrano GE, Beach TG, Bose S, Hansson O: The Relationship between p-tau217, p-tau231, and p-tau205 in the Human Brain Is Affected by the Cellular Environment and Alzheimer’s Disease Pathology. Cells 2024, 13.

46. Lilek J, Ajroud K, Feldman AZ, Krishnamachari S, Ghourchian S, Gefen T, Spencer CL, Kawles A, Mao Q, Tranovich JF, et al: Accumulation of pTau231 at the Postsynaptic Density in Early Alzheimer’s Disease. J Alzheimers Dis 2023, 92:241–260.

47. Jia S, Li Q, Rui X, Qin W, Zhang W, Dou J, Zhang X: The ubiquitin-proteasome system in Alzheimer’s disease: mechanism of action and current status of treatment. Front Aging Neurosci 2025, 17:1730206.

48. Tseng JH, Ajit A, Tabassum Z, Patel N, Tian X, Chen Y, Prevatte AW, Ling K, Rigo F, Meeker RB, et al: Tau seeds are subject to aberrant modifications resulting in distinct signatures. Cell Rep 2021, 35:109037.

49. Maniv I, Sarji M, Bdarneh A, Feldman A, Ankawa R, Koren E, Magid-Gold I, Reis N, Soteriou D, Salomon-Zimri S, et al: Altered ubiquitin signaling induces Alzheimer’s disease-like hallmarks in a three-dimensional human neural cell culture model. Nat Commun 2023, 14:5922.

50. Wang XJ, Yu J, Wong SH, Cheng AS, Chan FK, Ng SS, Cho CH, Sung JJ, Wu WK: A novel crosstalk between two major protein degradation systems: regulation of proteasomal activity by autophagy. Autophagy 2013, 9:1500–1508.

51. Lezoualc’h F, Engert S, Berning B, Behl C: Corticotropin-releasing hormone-mediated neuroprotection against oxidative stress is associated with the increased release of non-amyloidogenic amyloid beta precursor protein and with the suppression of nuclear factor-kappaB. Mol Endocrinol 2000, 14:147–159.

52. Pedersen WA, McCullers D, Culmsee C, Haughey NJ, Herman JP, Mattson MP: Corticotropin-releasing hormone protects neurons against insults relevant to the pathogenesis of Alzheimer’s disease. Neurobiol Dis 2001, 8:492–503.

53. Kim Y, Kim J, Son M, Lee J, Yeo I, Choi KY, Kim H, Kim BC, Lee KH, Kim Y: Plasma protein biomarker model for screening Alzheimer disease using multiple reaction monitoring-mass spectrometry. Sci Rep 2022, 12:1282.

54. Askenazi M, Kavanagh T, Pires G, Ueberheide B, Wisniewski T, Drummond E: Compilation of reported protein changes in the brain in Alzheimer’s disease. Nat Commun 2023, 14:4466.

55. Stepler KE, Mahoney ER, Kofler J, Hohman TJ, Lopez OL, Robinson RAS: Inclusion of African American/Black adults in a pilot brain proteomics study of Alzheimer’s disease. Neurobiol Dis 2020, 146:105129.

56. Drummond E, Pires G, MacMurray C, Askenazi M, Nayak S, Bourdon M, Safar J, Ueberheide B, Wisniewski T: Phosphorylated tau interactome in the human Alzheimer’s disease brain. Brain 2020, 143:2803–2817.

