## Supplemental Figures for "Targeted proteomics of postmortem human brain reveals neurobiologic heterogeneity in Alzheimer’s disease using NULISA technology"

**Contents**

**Supplementary Figures**

**Supplemental figure S1. Scatter plot of protein targets associated with age at onset**

**Supplemental figure S2. Scatter plot of protein targets associated with corticolimbic index (CLix)**

Supplemental Figure S1.

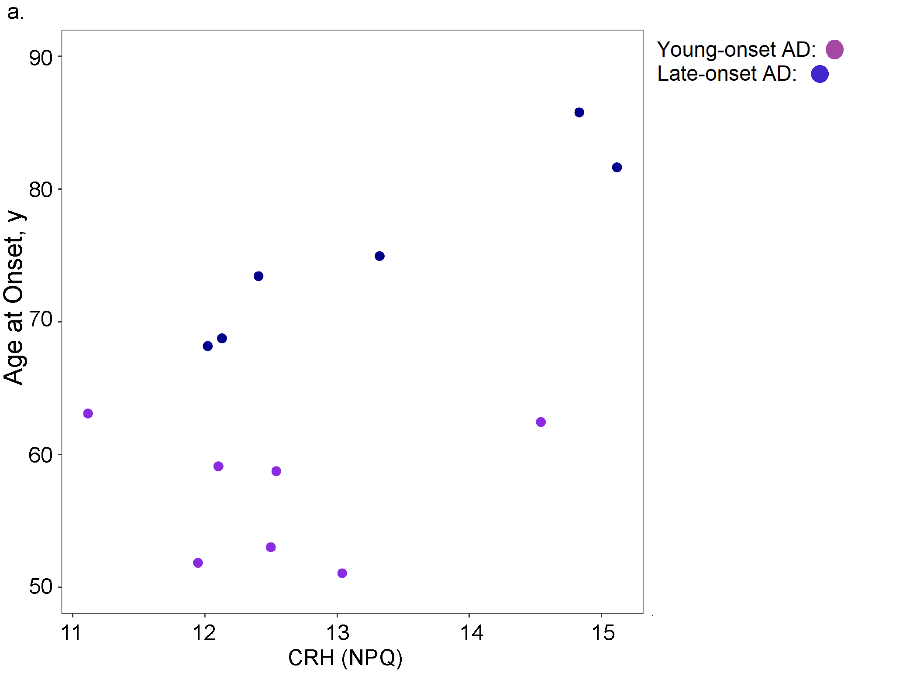

Supplemental figure S1. Scatter plot of CRH protein level plotted against age at onset. Young-onset AD cases are plotted in fuschia and late onset AD cases shown in violet. Abbreviations: CRH= corticotropin releasing hormone, NPQ= NULISA protein quantification unit of measure, young-onset= age at onset <65 years.

Supplemental Figure S2.

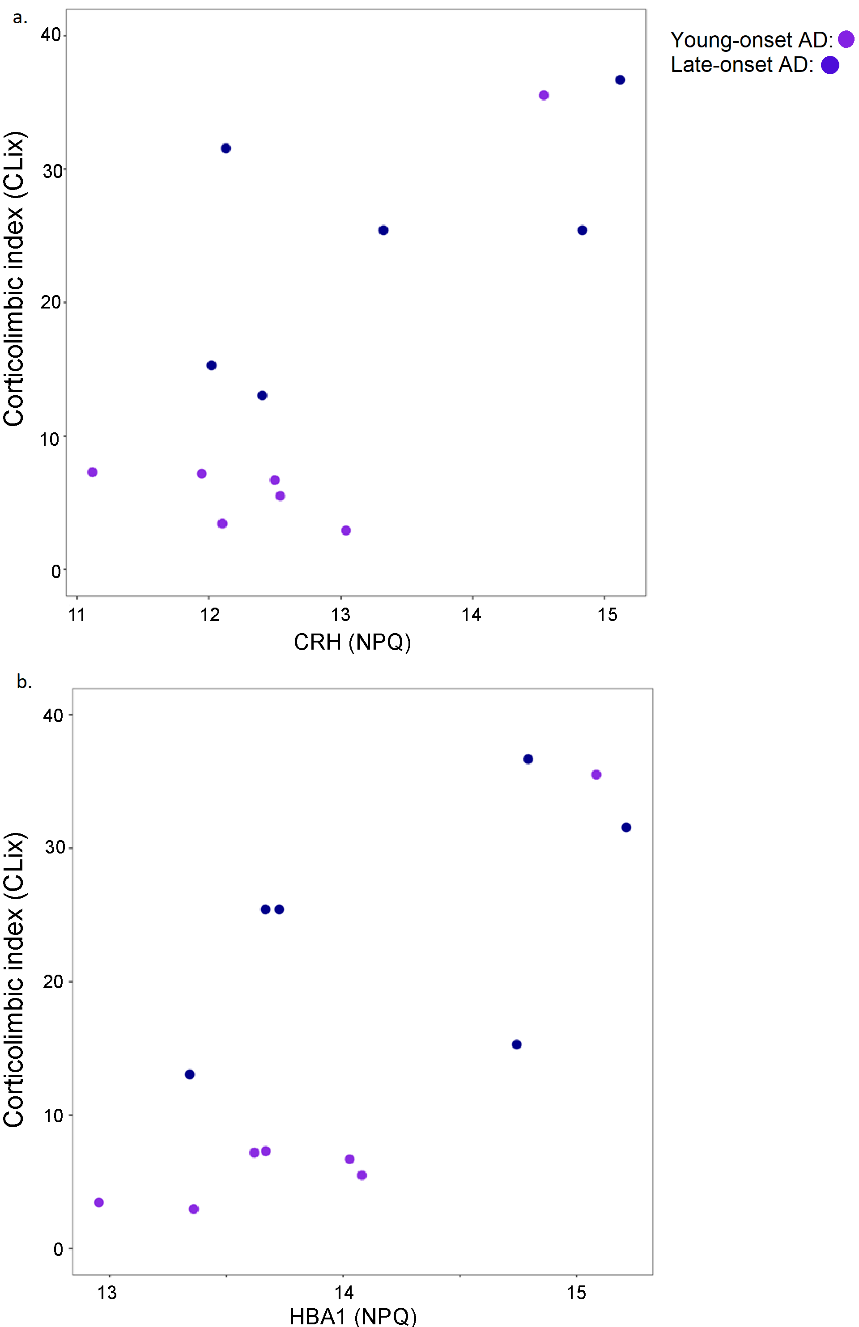
Supplemental figure S2. Levels of CRH and HBA1 are lower in cases with greater cortical tangle pathology. Scatter plots reflect protein levels of corticotropin releasing hormone (a) and hemaglobin alpha subunit-1 (b) plotted by corticolimbic index (CLix) score. Young-onset AD cases are plotted in fuschia and late onset cases plotted in violet. Abbreviations: CRH= corticotropin releasing hormone, HBA1= hemaglobin alpha subunit-1, NPQ= NULISA protein quantification unit, young-onset AD= age at onset <65 years.
